# Rule-based sequences in sooty mangabey vocal communication

**DOI:** 10.1101/2025.07.07.663462

**Authors:** Auriane Le Floch, Cédric Girard-Buttoz, Tanit Souha Azaiez, Natacha Bande, Roman M. Wittig, Steven Moran, Klaus Zuberbühler, Catherine Crockford

**Author notes:** Corresponding authors: Auriane Le Floch, ORCID iD 0000-0003- 3355-5830; Catherine Crockford. Joint senior authors (Klaus Zuberbühler, Catherine Crockford).

## Abstract

Investigating how non-human animals produce call sequences provides key insights into the evolutionary origins of meaning in vocal communication, including syntax. Many species combine calls into structured sequences, often following specific rules, yet most studies focus on only a few sequences per species. This limits our understanding of their ability to combine calls and convey meaning through sequences. Our study addresses this gap by documenting the vocal sequence repertoire and underlying rules of sooty mangabeys *(Cercocebus atys)*, a West African monkey species. We collected and analysed 1,672 recordings from two wild groups in the Taï National Park, Ivory Coast. Sooty mangabeys frequently combine calls but rely on a limited set of sequences. Within these, we identified rules of call ordering, recurrence and hierarchical structuring. Notably, they produced hierarchically structured sequences using only two call types, a previously unreported system in animal communication. Our findings suggest that sooty mangabeys use both structured and unstructured sequences, likely encoding different types of information. While context of production remains essential for determining meaning, our results highlight how a whole-repertoire approach reveals the full range of rule-based sequences and their potential for meaning expansion beyond the number of individual calls available in the repertoire.

## Introduction

One key feature of language is syntax, allowing humans to flexibly combine sounds into more complex utterances to generate a potentially infinite number of structure-meaning relations. The evolution of this ability, presumably unique to humans, is challenging to reconstruct ^1^. One powerful way to explore the problem is to study vocal signals in non-human animals ^2,3^ and how combining them into sequences can convey meaning. Relevant here is that, although humans are capable of producing and perceiving hundreds of phonemes (i.e., smallest units of sound in a language), all languages only utilise a fraction of them, with some languages using as few as 11 phonemes ^4,5^. This demonstrates that a small number of sounds is sufficient to sustain a language. The evolution of a communicative system capable of conveying a substantial range of meanings is thus likely less about producing increasingly rich vocal repertoires and more about evolving the ability to combine sounds in structured ways. This study investigates the vocal sequence repertoire of a *catarrhine* monkey, focusing on the rules that govern sequence structuring, independent of whether they convey specific meanings, to explore the potential for meaning generation through principles of sequence organisation.

Much research effort is currently being devoted to understanding the early origins of syntax and how it is integrated into the semantic system ^6^. This has led to a growing body of literature documenting how various species—including primates, birds, mongooses, marine mammals, elephants, hyraxes, bats, and amphibians—combine acoustically discrete sound units into larger sequences (frogs Bhat et al., 2022; elephants Hedwig & Kohlberg, 2024; see Engesser & Townsend, 2019 for a review on other species). Although not all sequences convey changes in meaning—typically songs, which are rarely meaning-specific and therefore outside the scope of this study ^9^ —recent research reveals that some sequences do, spanning a diverse range of species e.g., apes ^10–18^, bats ^19^, birds ^20–29^, elephants ^8^, frogs ^7^, mongooses ^30,31^, monkeys ^32–46^.

However, due to the time-consuming nature of this work, typically conducted through playback experiments (see Berthet et al., 2022 for review), only few such sequences have been reported per species. A promising approach to address this limitation is to first analyse entire vocal repertoires, focusing on whether sequences show rule-based organisation, irrespective of their semantic associations. One key metric in such analyses is flexibility—the extent to which calls in a vocal repertoire can be combined freely to form sequences ^48^. Flexibility can increase by expanding both the length of sequences and their diversity, i.e., the number of distinct sequences uttered. Diversity can also arise from adhering to specific rules that define how sequences in a vocal system are typically organised, including patterns of reoccurrence, ordering and hierarchical structures, where certain calls or strings of calls can be embedded within others, much like grammar in human language (see Berthet et al., 2022 for review and Table 1). Overall, few studies have investigated the natural vocal production of rule-based sequences in animals at the level of the entire repertoire, and even fewer have explored this in relation to associated meaning (see Table 1). Assessing vocal repertoire flexibility and diversity, therefore, provides a critical lens to further explore whether sequence structure may generate new meaning or simply reflect organisational variation without semantic implications.

**Table 1.** Rules in animal vocal sequences References for animal species were selected based on their relevance to meaning, except for hierarchical structures. For hierarchical structures, we included all species found in the literature, regardless of whether meaning was explicitly investigated in these studies. Songs were not included, as, although they can be hierarchically structured and offer interesting parallels with phonology, they are not typically meaning-specific ^9^. Note that, while classified into different categories, these rules are not necessarily mutually exclusive.

| Category | Rule | Description | Animal species |
| --- | --- | --- | --- |
| Reoccurrence | Repetition | Consecutive occurrence of the same call type at least two times, such as in the pattern ABBB, where the call 'A' is followed by multiple repetitions of 'B'. | Humayun's night frogs<sup>7</sup>, bonobos<sup>11</sup>, Carolina chickadees<sup>24</sup>, great tits<sup>26</sup>, pied babblers<sup>22</sup>, black-and-white colobus<sup>49</sup>, putty-nosed monkeys<sup>35</sup>, Thomas langurs<sup>45</sup>, indris<sup>44</sup> |
|  | Iteration | Repeated occurrence of a call type within a sequence, interspersed with other call types, such as in the pattern ABA, where 'A' is repeated with 'B' in between. | Bonobos<sup>11</sup> |
|  | Proportion | The overall number of times a call type appears in a sequence compared to the occurrence of one or other call types. | Amboli bush frogs<sup>7</sup>, titi monkeys<sup>36</sup> |
| Ordering | Positional bias | The tendency for a particular call type to occur at a specific position within a sequence. | Chestnut-crowned babblers<sup>20</sup>, Campbell's monkeys<sup>46</sup> |
|  | Gradation | Call types are arranged in a particular order within a sequence, forming a pattern that transitions from one type to another. | Meerkats<sup>31</sup>, black-fronted titi monkeys<sup>41</sup> |
| Hierarchical structure | Recursion | Elements are embedded within other elements of the same category<sup>50</sup>.<br>In animal vocal production, this means that call types are combined to create a pattern that is repeated within itself. For example, a simple sequence (AB) can expand recursively into ABABCAB, where part of the original pattern (AB) is embedded within itself. | Orangutans<sup>51</sup> |
|  | Recombination | Independently emitted short sequences are combined into longer sequences (e.g., AB becomes ABC). | Chimpanzees <sup>48</sup>, common marmosets <sup>52</sup> |
|  | Non-adjacent dependencies | A relationship between two elements that are separated in time or space and that cannot be explained only by the occurrence of multiple pairs of adjacent relationships <sup>53</sup>. For instance, a rule of non-adjacent dependencies would be that a sequence starting with 'A' must end with 'C', regardless of call types occurring in between (e.g., ABAC or ABBC). | Common marmosets <sup>52</sup> |

**Table 2.** Occurrence of calls used singly (single calls) and in sequences (combined calls) for each call type and sex. The columns % combined indicate the percentage of calls for calls produced in sequence only (i.e., excluding calls produced singly) for each call type and sex.

| Call type | Female |  |  | Male |  |  |
| --- | --- | --- | --- | --- | --- | --- |
|  | N single calls | N combined calls | % combined | N single calls | N combined calls | % combined |
| grunt | 968 | 391 | 42.4 | 225 | 6 | 2.3 |
| twitter | 133 | 342 | 37.1 | - | - | - |
| shrill | 35 | 18 | 2 | - | 101 | 38.7 |
| hoo | - | 48 | 5.2 | 1 | 103 | 39.5 |
| growl | 69 | 73 | 7.9 | 57 | 34 | 13 |
| scream | 49 | 34 | 3.7 | - | 8 | 3.1 |
| grumble | - | 16 | 1.7 | - | 5 | 1.9 |
| wau | - | - | - | - | 4 | 1.5 |
| vibrato | 110 | - | - | - | - | - |
| whoopgobble | - | - | - | 4 | - | - |

The main hypothesis for explaining species differences in flexibility is the amount of information that a species needs to convey ^54^. For this reason, many animal communication studies are concerned with the impact of structure on meaning, such as whether the meaning of a sequence is derived or distinct from the meaning of its constituent parts, e.g., apes ^12–14,16,17^, birds ^21,24^, monkeys ^34,46^, elephants ^8^, mongooses ^30^. Several distinct patterns have been found across the species that have been analysed. First, in some species one call type functions as a kind of affix that can be merged with an otherwise stand-alone call (e.g., ‘A’ becomes Ab), which then modifies the initial meaning, as observed in Campbell’s monkeys ^38,42^, mongooses ^55^ and Diana monkeys ^37^. Another mechanism is the production of distinct structures, i.e., repeating a specific call in a sequence, as in bonobos ^11^ or titi monkeys ^36^, or producing different call orders depending on event type, for example in meerkats ^31^, titi monkeys ^41^, Chestnut-crowned babblers ^20^ and Campbell’s monkeys ^46^.

Understanding the evolutionary drivers and constraints of flexibility, therefore, is likely to advance our understanding of what exactly led to language in humans. This requires a systematic, whole-repertoire approach, which involves assessing all possible sequences within a repertoire along with their flexibility, including the potential rules that govern their organisation—an approach that has rarely been explored, e.g., chimpanzees ^15,48^, marmosets ^52^. Such an approach is a critical first step to assess the potential of a given repertoire to expand meanings through call combinations. Here, we addressed this gap with an observational study of wild sooty mangabey vocal behaviour, a primate with a call repertoire comparable in size to great apes ^56,57^ and suggested containing several call sequences ^57,58^. We were particularly interested in identifying which calls from the repertoire were combined, how long and diverse the resulting sequences were, and whether it was possible to discern rules in their composition.

## Methods

### Data collection

ALF, NB and TSA collected data on two groups of sooty mangabeys habituated to human presence, the Taï Monkey Project (TMP) group from May to July 2022 and the Taï Chimpanzee Project (TCP) group from February to October 2023 at the Taï National Park (Ivory Coast). We used half-day continuous focal animal sampling ^59^ on adult individuals from dawn to midday or from midday to dusk. We recorded all the calls produced by the focal individuals on an all-occurrence basis, as well as calls produced by known adult individuals in visual range on an ad-libitum basis, using directional microphones (Sennheiser models K6, MK 316, MKH 416 P48 and MKH 416T-F) and recorders (Marantz Solid State models PMD661 and PMD661 MKII), digitised at a 44,1 kHz sampling rate and 16-bit sampling depth in a ‘.wav’ format.

This work was conducted under the research permit for the Taï Chimpanzee Project (Wittig/006/MESRS/DRI) from the Ivorian Ministry of Higher Education and Scientific Research and the Ivorian Office of Parks and Reserves for access to the Taï National Park. This study was purely observational with audio recordings and as such approved by the “Ethikrat der Max Planck Gesellschaft”. The use of wild animals in this research adheres to the guidelines set forth by the Association for the Study of Animal Behaviour and the International Union for Conservation of Nature.

### Audio recording coding

ALF, NB and TSA coded audio recordings using Raven Pro (ALF v1.6.5; NB and TSA v1.6.4)^60^. We visually inspected spectrograms (FFT size 1024, Hann window, hope size 256 samples) while listening to the recordings to identify and manually annotate each sound element with start and end times (see glossary for definition of an element). We used sooty mangabey vocal repertoire ^57^ and single out 10 call types: ‘grunt’, ‘twitter’, ‘growl’, ‘scream’, ‘grumble’, ‘vibrato’ (referred to as ‘copulation call’ in Range & Fischer’s vocal repertoire), ‘whoop-gobble’, ‘wau’, ‘shrill’ (the first high-frequency sound elements of the alarm call in Range & Fischer’s vocal repertoire) and ‘hoo’ (referring to both the ‘hoo’ call described in Range & Fischer’s vocal repertoire and the last, optional low-frequency sound element that was initially described as part of the alarm call). All call types used in this study are detailed in the supplementary materials together in the section Document S1 with spectrographic examples and corresponding definitions from Range & Fischer’s vocal repertoire.

To define a call, we applied a 2-second threshold for intervals between elements of the same type. This rule was chosen to avoid overinflating the number of calls by splitting continuous vocalisations. Of the 8,259 intervals measured across 1,693 utterances, 98.18% were shorter than 2 seconds, with a median of 0.09 seconds and a standard deviation of 1.82 seconds. By using this threshold, we ensured that only intervals longer than 2 seconds were treated as breaks between distinct calls, while the majority of shorter intervals remained within the same call.

To define a sequence, we set a 1-second maximum interval threshold based on the distribution of 1,122 intervals between distinct calls, measured across 619 vocal utterances. The majority of intervals were short, with a median of 0.27 seconds, and 69.4% of intervals fell below 1 second. The standard deviation was 6.39 seconds, reflecting considerable variability. The 1- second threshold was chosen to be conservative, capturing typical call intervals while excluding longer breaks, ensuring that sequences were not overinflated by including excessive pauses.

### Glossary

| Term | Definition |
| --- | --- |
| Vocal element | A continuous sound with a distinct beginning and end. In rare cases, two different types of elements may occur without a clear silent gap between them; in such cases, they are distinguished using their specific acoustic properties <sup>61</sup> (see supplementary materials for details on acoustic properties of elements). |
| Call type / Vocal unit | A call consisting of one or multiple elements of the same type, produced in succession with intervals of less than two seconds between them. |
| Vocal sequence | A series of different consecutive call types (or vocal units), where each call type follows the previous one with an interval of less than one second between them. |
| X-call type sequence | A vocal sequence containing X distinct call types, which may or may not include one or more iterations. For example, a two-call type sequence can be AB, ABA, or ABABA. |
| Sequence length | The number of call types in a vocal sequence was measured in two ways: including iterations (e.g., ABA) to capture the raw sequence length, and excluding iterations (e.g., ABA becomes AB) to determine how many different call types are combined. |
| Combination pair | Two call types produced consecutively within a sequence, regardless of order. For example: the sequence ABC contains two pairs: A and B, and B and C; the sequences AB and BA hold the same combination pair: A and B. |
| Sequence diversity | The number of unique sequences, defined by the specific order of call types. With iterations considered, sequences like AB and ABA are unique. Without iterations, a unique sequence (e.g., AB) includes all sequences starting with AB, such as AB, ABA, and ABAB. |
| Vocal utterance | Either a single call type (vocal unit) produced in isolation (not part of a sequence) or a vocal sequence. |

### Analyses

We first visually inspected the sequence flexibility by looking at (1) the number of different call types in the repertoire that can be combined, (2) the sequence length with and without iterations, (3) the number of different possible combination pairs and (4) the sequence diversity with and without iterations (see glossary for definitions). We then focused on a subset of the data for which we had sufficient sample size to analyse their rules: ordering was investigated on 2-call type sequences (93% of the sequence production) and reoccurrence on sequences composed of ‘grunts’ and ‘twitter’ (61% of the sequences, see Tables S1, S2, S3 and S4 for details). Finally, still focusing on sequences of ‘grunts’ and ‘twitters’, we assessed their hierarchical structures based on their composition and rules.

### Ordering

To analyse ordering in 2-call type sequences, we examined positional bias based on the first occurrence of each call type (i.e., whether they were more likely to occur in the first or second position within the sequence). 3-call type and 4-call type sequences constituted only 7% of the total sequences produced (*N*=26 sequences) and were excluded from the analysis due to insufficient sample size (Figure 3, see Tables S1 and S3 for details). We investigated six call types (‘shrill’, ‘grunt’, ‘growl’, ‘scream’, ‘twitter’ and ‘hoo’). However, analyses were not conducted for ‘shrill’ and ‘hoo’ because these call types were consistently produced in the same positions (first and second, respectively). The remaining call type appearing in 2-call type sequence, ‘grumble’, was excluded due to low sample size (*N*=4). We used sequences produced by both sexes for the analysis. Bayesian logistic regression models were employed to examine call position (first/second) as a binary outcome variable. This resulted in four models being constructed for the following call types: ‘grunt’, ‘growl’, ‘scream’ and ‘twitter’. These models included the group (TMP/TCP) as fixed effect (for models with at least five sequences sampled from each group: model ‘twitter’, model ‘grunt’, model ‘growl’, model ‘scream’) and the identity of the caller as random intercept (for models with at least 50% of the callers sampled that produced at least two sequences: models ‘twitter’ and ‘grunt’) to account for repeated samples of the same individual in the same group.

### Reoccurrence

We started by analysing iterations in sequences of ‘grunts’ and ‘twitters’ (44.5% of the ‘grunt’ and ‘twitter’ sequences). Specifically, we aimed to identify which call type (‘grunt’ or ‘twitter’) was more likely to occupy the final position, depending on which call type initiated the sequence. We first used a Bayesian logistic regression model to analyse the type of last call (‘grunt’/’twitter’) as binary outcome variable for sequences starting with a ‘grunt’. We included the group (TMP/TCP) as a fixed effect and the identity of the caller as a random intercept to account for repeated samples of the same individual in the same group. We then ran another model, analogous to the previous model on sequences starting with ‘grunt’, but this time with sequences starting with ‘twitter’. This model was built almost identically, except that we did not include the group (TMP/TCP) as fixed effect because it had less than five sequences sampled from the TMP group.

We then examined the repetitions of elements within calls in sequences of ‘grunts’ and ‘twitters’, depending on the call position within the sequence. For this analysis, we used sequences both with and without iterations. We excluded sequences of certain lengths if they were produced by fewer than five individuals to ensure a sufficiently representative sample size for each sequence length. We used Bayesian Poisson regression models to analyse the number of repetitions of elements in each call as a function of their position within the sequence: first, middle, or last call. Separate models were run for each sequence and call type. Sequence types were defined by whether they started with a ‘grunt’ or a ‘twitter’. Call types were ‘grunt’ and ‘twitter’. For ‘grunts’, we examined the first ‘grunt’ call, the middle ‘grunt’ call(s) (i.e., all calls occurring between the first and last ‘grunt’ calls in the sequence, intercepted by ‘twitter’ calls), and the last ‘grunt’ call. For example, in the sequence grunt_twitter_grunt_twitter_grunt, there is one middle ‘grunt’ call in between the two ‘twitters’, but in a sequence like grunt_twitter, there is only one first ‘grunt’ call, no middle or last ‘grunt’ call. The same analysis was applied to ‘twitters’, considering the first, middle, and last ‘twitter’ calls within sequences. We added the length of the sequence and the group (TMP/TCP) as fixed factors and accounted for individual variability through a random intercept. The first model looked at the repetitions of ‘grunt’ elements in grunt_twitter sequences (i.e., sequences starting with a ‘grunt’) depending on the position of each ‘grunt’ call: first ‘grunt’ call appearing in the sequence, middle and last ‘grunt’ calls. The second model looked at the repetitions of ‘twitter’ elements in those same grunt_twitter sequences depending on the position of each ‘twitter’ call. After fitting these two models, we performed pairwise comparisons between the levels of the variable position (first/middle/last) using the posterior distribution of the estimated number of repetitions for each position (i.e., estimated marginal means approach). Then, analogous to the two first models on grunt_twitter sequences, we ran our two other models to look for repetitions of elements in twitter_grunt sequences (i.e., sequences starting with ‘twitter’) of ‘grunt’ and ‘twitter’ calls, respectively.

### Model specifications

For all our models, we used the default flat priors from the *brms* package version 2.20.4 ^62^: the intercept had a weakly informative normal prior centered at 0 with a standard deviation of 10, the fixed effect for the predictor (group) a normal prior centered at 0 with a standard deviation of 2.5, and the standard deviation of the random intercepts and a Student-t prior with 3 degrees of freedom, centered at 0, and with a scale parameter of 10). We ran the models using four Markov chains, each with 4,000 iterations, including a warm-up of 2,000 iterations per chain. This resulted in a total of 8,000 post-warm-up samples. We manually set the *adapt delta* parameter either to 0.95 (positional bias models ‘twitter’, model ‘grunt’) or 0.90 (positional bias models ‘growl’ and ‘scream’ and the four models on element repetitions) ensuring thorough exploration of the posterior distribution and minimising the likelihood of divergent transitions, except for the models call iterations in grunt_twitter sequences and call iterations in twitter_grunt sequences where we kept the default (0.80). Convergence of the Markov chains was evaluated using the potential scale reduction factor R-hat, with values less than 1 indicating adequate convergence. We also visually inspected trace plots for each parameter to ensure stable and well-mixed chains. Posterior predictive checks were conducted to assess the fit of the models to the data.

The analyses were conducted using R version 4.3.3 ^63^ with the *brms* package version 2.20.4 ^62^. The comparisons were conducted using the *emmeans* package version 1.10.0 ^64^.

### Hierarchical structures

We used the rules of ordering and reoccurrence we identified to write down the sequences composed of ‘grunts’ and ‘twitters’ and their hierarchical structures in a table, similar to writing down the grammar of a language.

## Results

We conducted continuous focal recordings on adult sooty mangabeys resulting in TCP females: *N*=17 individuals, *N*=232h and 27min (mean per individual = 13h and 40min, SE=0h and 56min); TMP females: *N*=18 individuals, *N*=262 hours and 45 minutes (mean per individual = 14h and 35min, SE=3h and 28min); TCP males: *N*=8 individuals, *N*=122h and 21min (mean per individual = 15h and 17min, SE=1h and 39min); TMP males: *N*=4 individuals, *N*=10h and 30min (mean per individual = 2h and 37min, SE=0h and 24min).We also recorded vocal utterances from adlib individuals (i.e., who were not focal individuals): *N*=4 TCP females; *N*=2 TMP females; *N*=5 TMP males. From this, we collected and analysed a total of *N*=1,672 recordings comprising *N*=2,031 vocal utterances, the majority consisting of only one call type (females: *N*=1,364, 82.9% of all utterances; males: *N*=287, 75.9% of all utterances; Figure 3).

### Call types

Females used seven of eight call types in sequences. Among these, ‘grunts’ and ‘twitters’ were the most frequently found in sequences, accounting for 42.4% and 37.1% of all sequences, respectively (Figure 1). These were followed by ‘growls’ (7.9%), ‘hoos’ (5.2%), ‘screams’ (3.7%), ‘shrills’ (2%) and ‘grumbles’ (1.7%).

**Figure 1.**
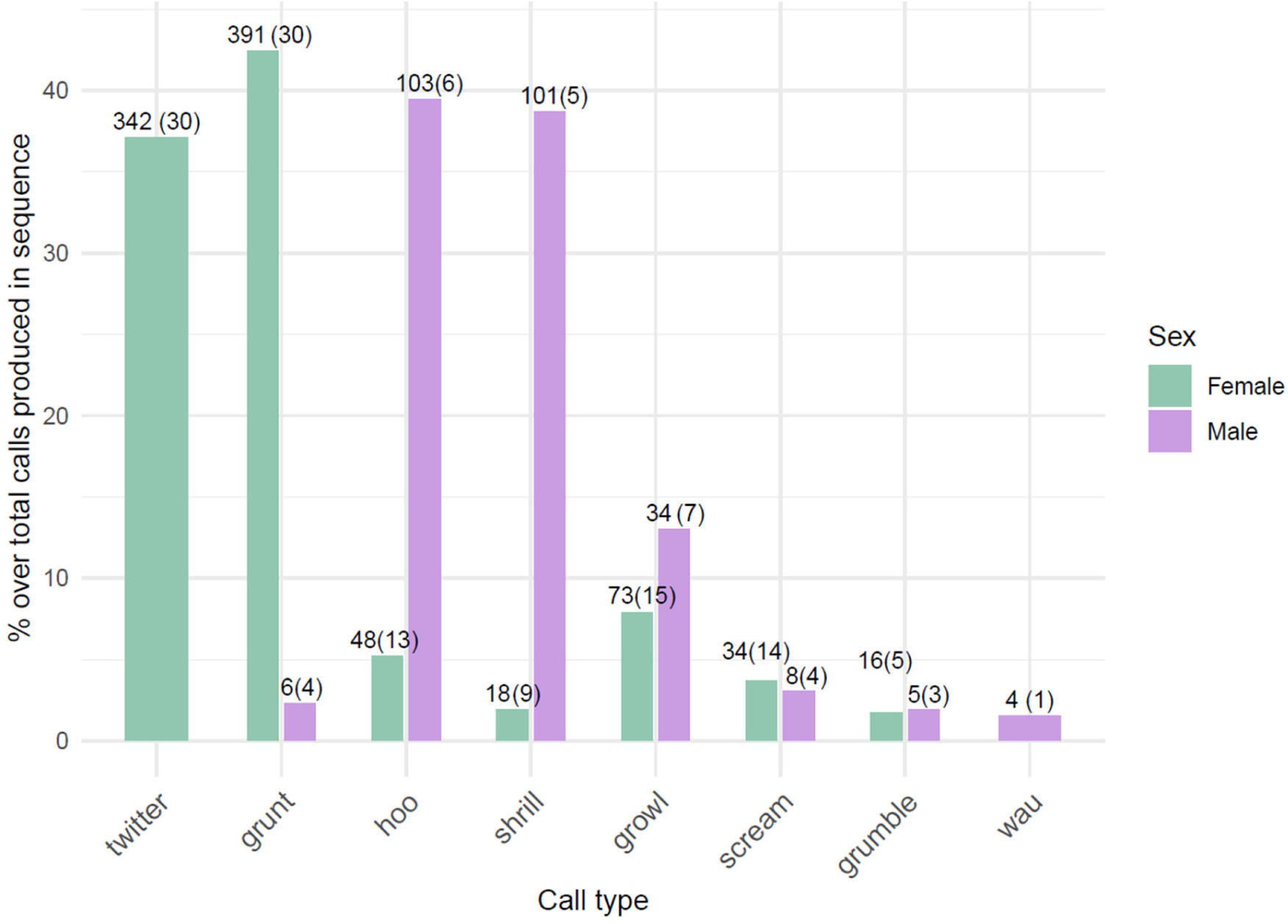
Frequency of call types used in sequences. These numbers represent all occurrences of calls found in sequence, i.e., including iterations. Above each bar is indicated the sample size with the number of calls recorded next to the number of individuals producing them in brackets.

Males also used seven of eight call types in sequences, with a large dominance of ‘hoos’ (39.5%) and ‘shrills’ (38.7%). Other, less frequently combined call types were ‘growls’ (13%), ‘screams’ (3.1%), ‘grunts’ (2.3%), ‘grumbles’ (1.9%) and ‘waus’ (1.5%).

Although both sexes combined a similar number of call types into sequences, females used most of their calls both singly and in sequence, with only two of seven call types exclusively used in sequences. In contrast, males primarily used calls in sequence, with five of seven call types used exclusively in sequences (Figure 2). Note that while one ‘hoo’ call was produced singly, this occurred only once, so we classified this call as being produced exclusively in sequence.

**Figure 2.**
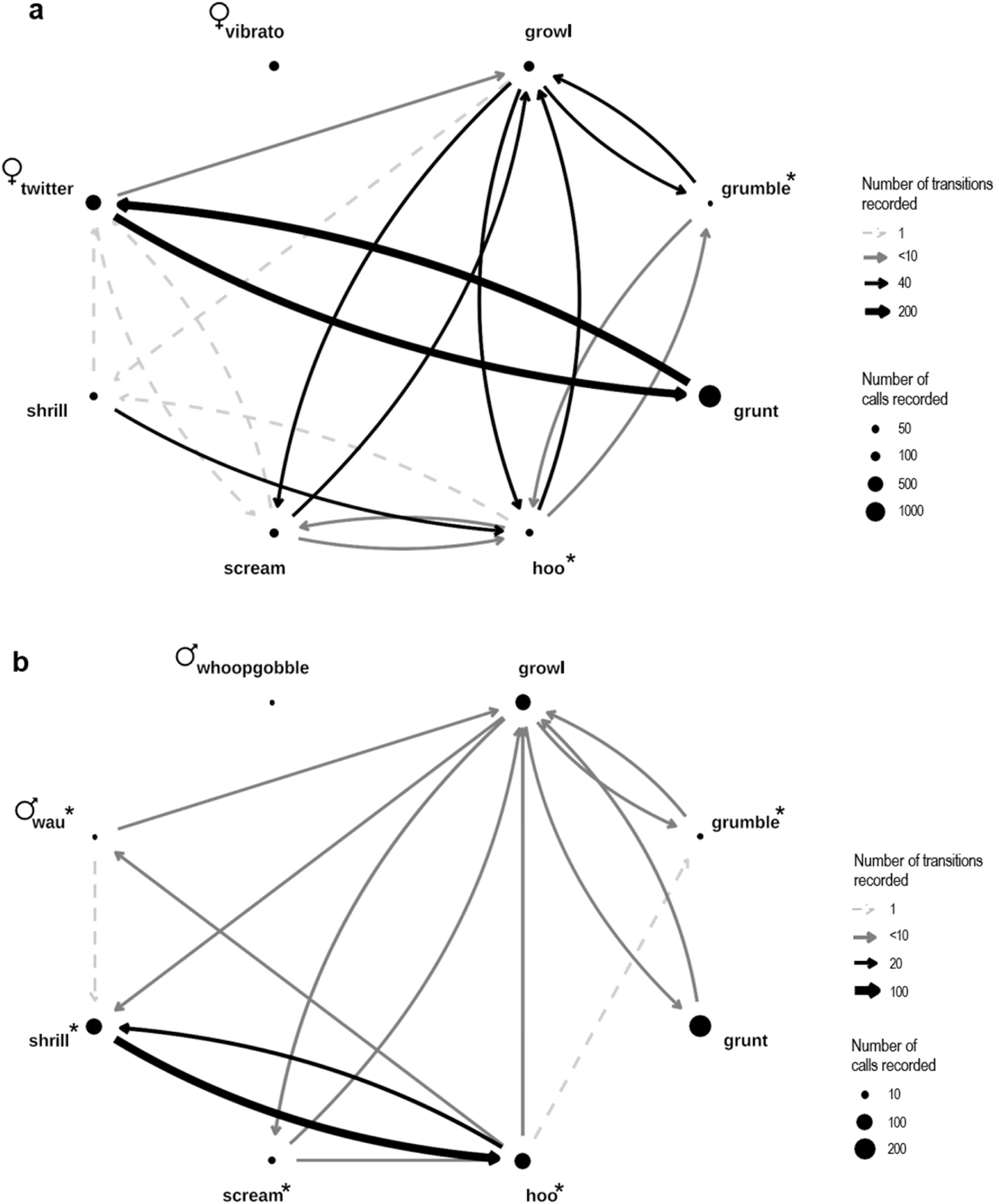
Vocal repertoire of adult sooty mangabey a females and b males. These networks represent all recorded call types and transitions between the different call types in sequences, including iterations. * = call type produced exclusively in sequence with another call type (i.e., never or idiosyncratic -maximum one occurrence- singly). **♀** = call type produced only by females. ♂ = call type produced only by males. Node (circle) size is proportional to the occurrence of call for each call type and edge (line) thickness and color are proportional to the number of recorded transitions. See Tables S2 and S4 for detailed sample sizes.

**Figure 3.**
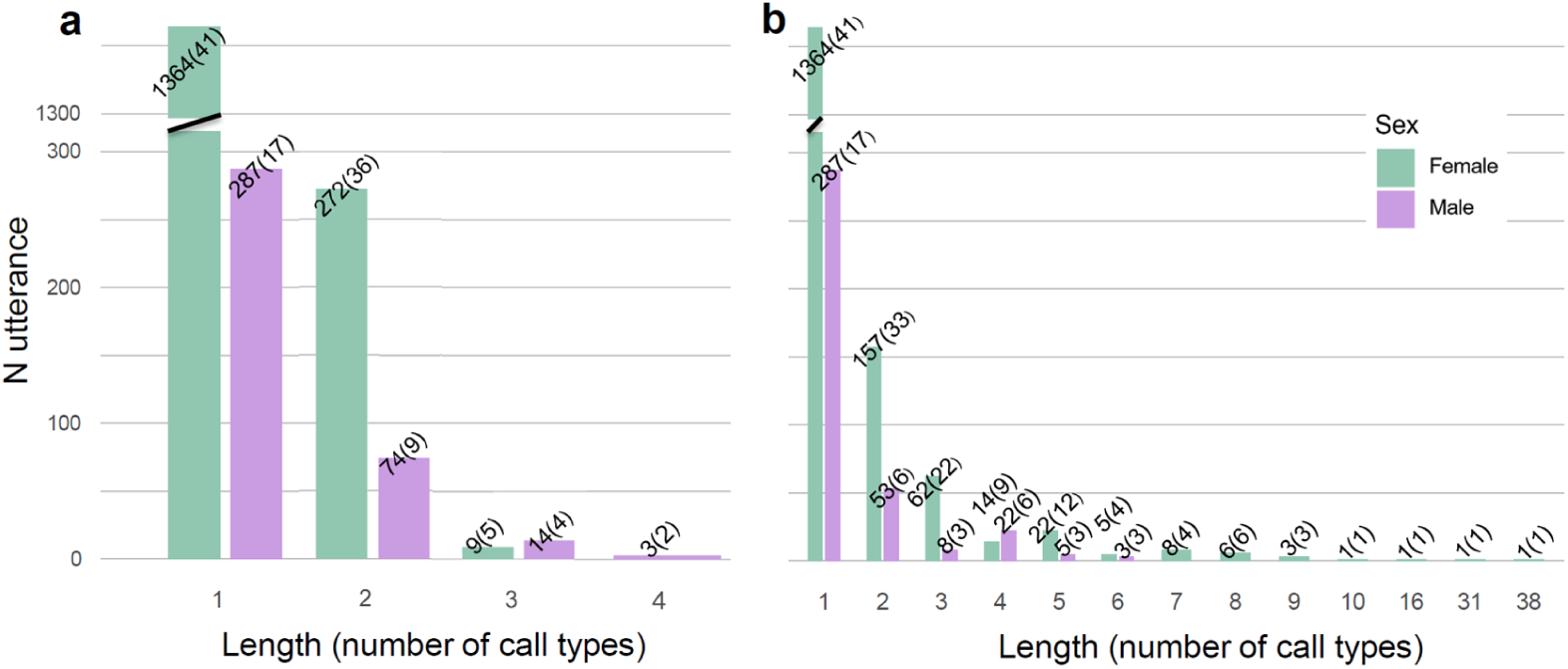
Number of utterances recorded per length and sex a without iteration and b with iteration Above each bar is indicated the sample size with the number of utterances recorded next to the number of individuals producing sequences of that length in brackets.

### Combination pairs

In females, most combination pairs (78.2%) were between ‘grunts’ and ‘twitters’, (Figure 2 and Figure S1). Combination pairs between ‘growls’ and ‘hoos’ accounted for another 7.2%, between ‘growls’ and ‘screams’ 4.8% and between ‘growls’ and ‘grumbles’ 3.4%. In males, most combination pairs were between ‘shrills’ and ‘hoos’ (71.2%). These were followed by combination pairs between ‘growls’ and ‘screams’ (6.5%), ‘growls’ and ‘hoos’ (5.3%), ‘growls’ and ‘grumbles’ (4.7%) and ‘growls’ and ‘grunts’ (4.1%) (Figure 2 and Figure S1).

### Sequence length

The most frequent sequences were composed of two call types for both sexes (Figure 3). Not counting iterations, the maximum sequence length consisted of four different call types for males (although these were infrequent; three sequences produced by two individuals) and three for females (Figure 3a). For sequences with iterations (i.e., repeated occurrences of one call type), the most frequent length was still two calls, for both sexes. In females, typical sequence length ranged up to nine calls, exceptionally consisting of 38 calls, though these were observed in single individuals. In males, sequences ranged up to six calls (Figure 3b).

### Sequence diversity

Without iterations, females produced 17 different sequences. The most frequent sequence was grunt_twitter (54.4%) followed by twitter_grunt (26.3%). With iterations, females produced 41 different sequences. Here again, the most frequent was grunt_twitter (29.2%) followed by grunt_twitter_grunt (15.7%) and twitter_grunt (15.7%).

Males, on the other hand, produced 11 unique sequences without iterations. The most frequent among these was shrill_hoo (72.5%) followed by shrill_hoo_growl (6.6%). When accounting for iterations, males produced 21 unique sequences. The most frequent sequence was shrill_hoo (53.8%) followed by shrill_hoo_shrill_hoo (17.6%). For further details, including sample size, please refer to Tables S1, S2, S3 and S4.

### Rules

#### Ordering - positional bias in 2-call type sequences

We found four call types with positional bias in sequence and two that did not have consistent bias (Table 3 and Figure 4). All ‘shrills’ were found in first position and all ‘hoos’ in second position. ‘Shrills’ only occurred in shrill_hoo sequences whereas ‘hoos’ could be combined, although less frequently, with other call types. We found consistent statistical support for ‘grunt’ to be biased towards the first position (Table 3 and Figure 4). Most ‘grunts’ were part of grunt_twitter and twitter_grunt sequences. We found some support, though with greater uncertainty, for ‘twitters’ to be biased toward the second position (Table 3 and Figure 4). Similar to ‘grunts’, most ‘twitters’ were found in grunt_twitter and twitter_grunt sequences. We did not find consistent support for a positional bias for ‘growls’ and ‘screams’ (Table 3 and Figure 4; see Table S5 for detailed results and Figure S2 for posterior predictive checks of the models).

**Figure 4.**
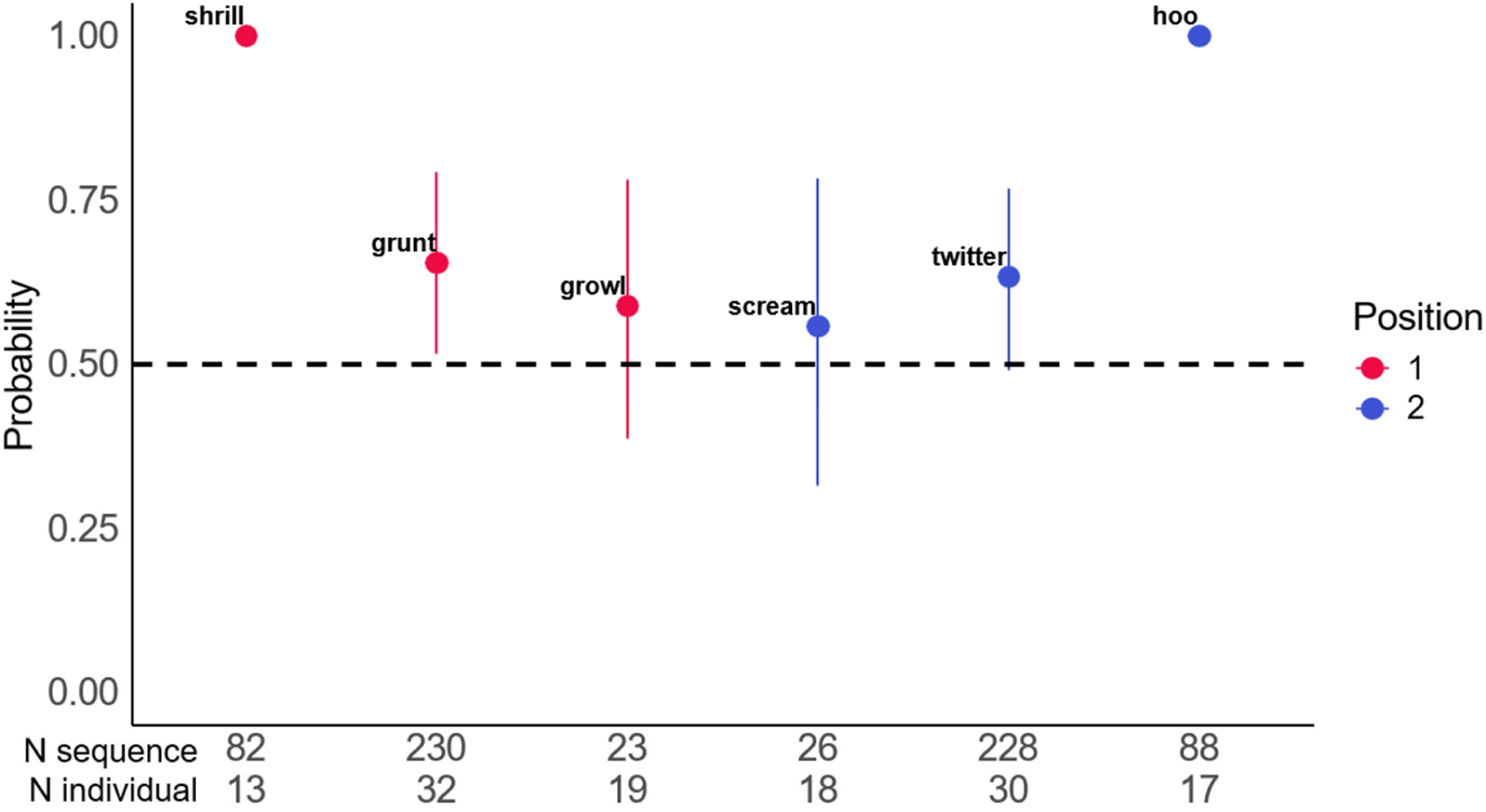
Probability of call types to occur in the first (red) or second (blue) position in 2-call type sequences The probability is calculated using either (1) the mean posterior distribution of each model (‘grunt’/’growl’/’scream’/’twitter’) or (2) the mean occurrence of the raw data (‘shrill’/’hoo’). The bars represent the 95% credible interval.

**Table 3.** Positional bias model results Call types with a 95% confidence interval (CI) range that does not overlap 0.5 indicate statistically significant positional bias. Sequence types are presented here without iterations so we understand which call type of the 2-call type sequences comes first and which comes second. Sequence types and sample sizes for ‘shrill’ and ‘hoo’ are included, although these were not analysed through models.

| Call type | Position | Mean posterior distribution | 95% CI range | Sequence type | N sequence |
| --- | --- | --- | --- | --- | --- |
| shrill | 1 | - | - | shrill_hoo | 82 |
| hoo | 2 | - | - | shrill_hoo<br>growl_hoo<br>scream_hoo | 82<br>3<br>3 |
| grunt | 1 | 0.65 | 0.52-0.79 | grunt_twitter<br>twitter_grunt<br>grunt_growl | 153<br>74<br>3 |
| twitter | 2 | 0.63 | 0.49-0.77 | grunt_twitter<br>twitter_grunt<br>twitter_growl | 153<br>74<br>1 |
| growl | 1 | 0.59 | 0.39-0.78 | growl_scream<br>scream_growl<br>growl_grumble<br>growl_hoo<br>grunt_growl<br>grumble_growl<br>twitter_growl | 16<br>7<br>3<br>3<br>3<br>1<br>1 |
| scream | 2 | 0.56 | 0.32-0.78 | growl_scream<br>scream_growl<br>scream_hoo | 16<br>7<br>3 |

<u>Reoccurrence - iterations of calls in sequences composed of ‘grunts’ and ‘twitters’</u> Considering sequences composed of ‘grunts’ and ‘twitters’ occurring in iteration and starting with a ‘grunt’, we found consistent support that the last call in a sequence was more likely a ‘grunt’ than a ‘twitter’ (mean probability and 95% CI for ‘grunt’ to be last 0.97, 95%-CI[0.88-1]). However, for sequences starting with ‘twitter’, we did not find that one call type was consistently more likely to be the last call than the other (mean probability and 95% CI for ‘grunt’ and ‘twitter’ to be last 0.65, 95%-CI[0.28-94] vs 0.35 95%-CI[0.06-72]) (Figure 5a; see Table S6 for detailed results and Figure S3 for posterior predictive checks of the models).

**Figure 5.**
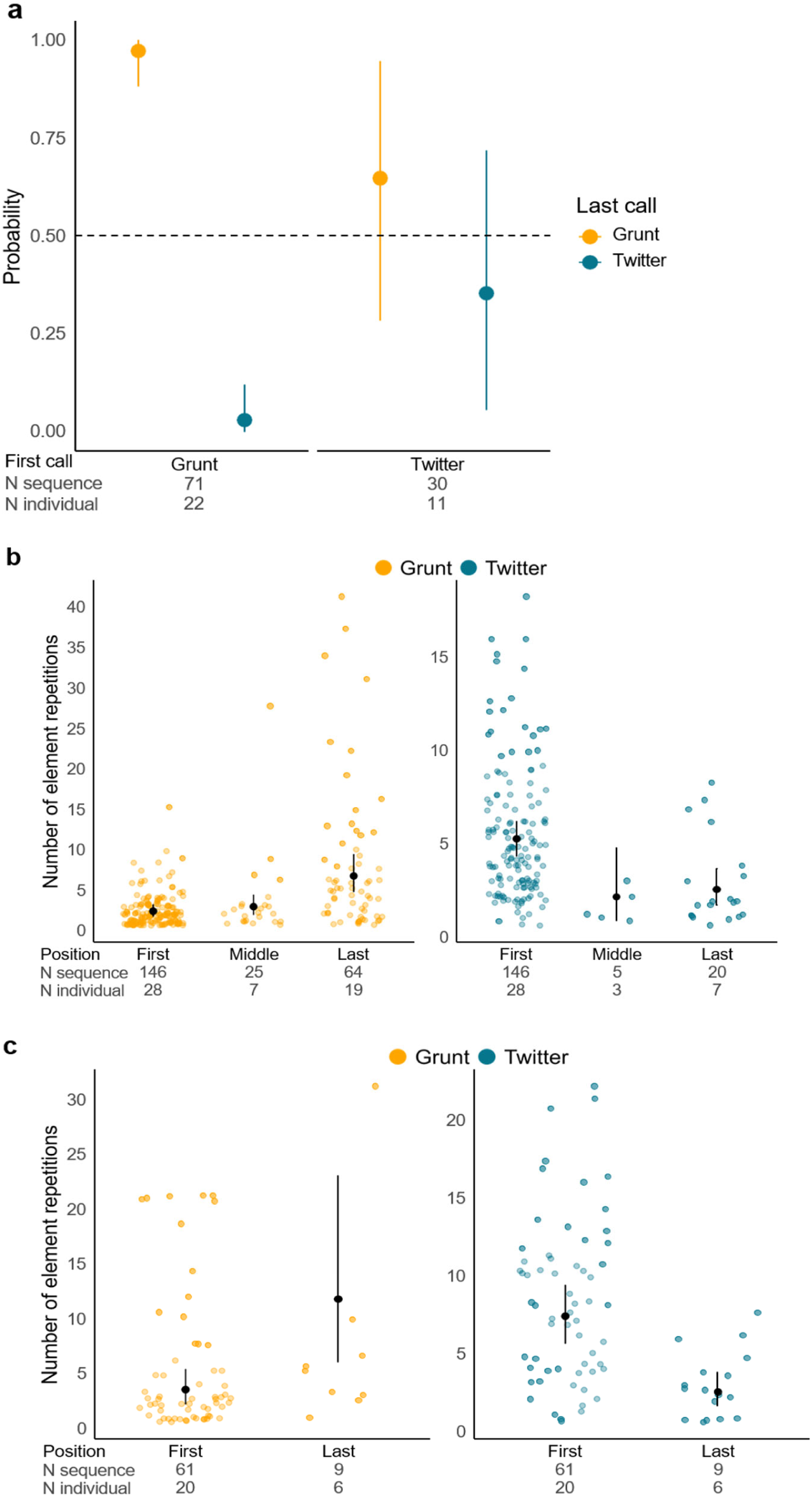
a Probability of ‘grunt’ (yellow) and ‘twitter’ (blue) to be produced as the last call type in iterative sequences composed of ‘grunts’ and ‘twitters’, and either starting with ‘grunt’ (left) or ‘twitter’ (right). b Number of element repetitions of ‘grunt’ calls (yellow/left) and ‘twitter’ calls (blue/right) depending on their position in sequences starting with a ‘grunt’. c Number of element repetitions of ‘grunt’ calls (yellow/left) and ‘twitter’ calls (blue/right) depending on their position in sequences starting with a ‘twitter’. The probability is calculated using the mean posterior distribution of each model. The bars represent the credible interval at 95% and the point is the mean posterior distribution.

Reoccurrence - repetition of elements within calls in sequences composed of ‘grunts’ and ‘twitters’

In sequences starting with a ‘grunt’, the last ‘grunt’ call contained more repetition of elements than the first and the middle ‘grunt’ calls (first ‘grunt’ calls mean posterior distribution = 2.40, 95%-CI[1.80-3.19]; middle ‘grunt’ calls mean posterior distribution = 2.96, 95%-CI[1.98-4.41]; last ‘grunt’ calls mean posterior distribution = 6.71, 95%-CI[4.78-9.37]). In these same sequences, we found consistent support for the first ‘twitter’ call to have more repetitions than the middle and the last ‘twitter’ calls (first ‘twitter’ calls mean posterior distribution = 5.29, 95%- CI[4.36-6.22]; middle ‘twitter’ calls mean posterior distribution = 2.17, 95%-CI[0.90-4.78]; last ‘twitter’ calls mean posterior distribution = 2.56, 95%-CI[1.74-3.68]). (Figure 5b).

The same pattern also applies to sequences starting with a ‘twitter’, with the last ‘grunt’ call having more element repetitions than the first ‘grunt’ call (first ‘grunt’ calls mean posterior distribution = 3.54, 95%-CI[2.25-5.35] vs last ‘grunt’ calls mean posterior distribution = 11.80, 95%-CI[6.10-23.06]) and the first ‘twitter’ call more repetition than the last one (first ‘twitter’ calls mean posterior distribution = 7.41, 95%-CI[5.67-9.39] vs last ‘twitter’ calls mean posterior distribution = 2.58, 95%-CI[1.70-3.83]). (Figure 5c; see Table S7 for detailed results and Figure S4 for posterior predictive checks of the models).

Hierarchical structures in sequences composed of ‘grunts’ and ‘twitters’

We found that sequences composed of ‘grunts’ and ‘twitters’ followed hierarchical structures of recombination, non-adjacent dependencies and recursion, as proposed in the Table 3.

## Discussion

This study assessed the potential for a *catarrhine* monkey vocal repertoire to produce new meaning through sequences. We quantified sooty mangabey sequence types and analysed their rules of ordering and recurrence, from which we identified hierarchical structures. We found that positional bias and rules governing call iterations and element repetitions within sequences included recombination, non-adjacent dependencies and recursion. Whether these rules change the meaning conveyed to receivers compared to when each call is emitted alone remains to be tested. We discuss these rules and compare the sooty mangabey flexibility in sequence production with that of other species.

### Rules and hierarchical structures

Structural patterns in the two dominant combination pairs—’shrills’ with ‘hoos’ and ‘grunts’ with ‘twitters’—suggest different mechanisms. ‘Shrills’ and ‘hoos’ showed strong positional bias, with ‘shrill’ first and ‘hoo’ second in 100% of sequences. ‘Shrills’ occurred alone, but not ‘hoos’ (one occurrence recorded). Similarly, sympatric-living Campbell’s monkeys add a suffix ’oo’ to an alarm call changing the meaning from a high-urgency to a lower-urgency threat ^38,42^. Both ‘shrills’ and ‘hoos’ are produced in alarm contexts ^57,65^, but confirming whether specific predator information is conveyed to conspecifics requires experimental studies. Sequences composed of ‘grunts’ and ‘twitters’ also showed strong positional bias, with ‘grunts’ being more likely in first position. Unlike shrill_hoo sequences, reverse sequences were also relatively prevalent, with ‘twitter’ in the first position in 32.6% of ‘grunt’ and ‘twitter’ sequences (Table 3).

When specifically analysing sequences composed of ‘grunts’ and ‘twitters’, additional rules of the reoccurrence of each call were evident. Sequences starting with a ‘grunt’ call more likely ended with a ‘grunt’ call, which was not the case for sequences starting with a ‘twitter’ call (Figure 5a). Moreover, regardless of which call started the sequence, the first ‘twitter’ call of the sequence had more element repetitions compared to ‘twitter’ calls produced later in the sequence, and likewise for the last ‘grunt’ call in a sequence (Figure 5b&c).

Also in ‘grunt’ and ‘twitter’ sequences, sooty mangabeys appear to produce hierarchical structures, using rule-based iteration and repetition indicating recombination, non-adjacent dependencies and recursion (Table 4). In contrast, marmosets and chimpanzees also use recombination (chimpanzees) and non-adjacent dependencies (marmosets) but with a broader range of call types ^48,52^. Moreover, hierarchical structures are rarely described in animal communication outside of song, with only orangutangs being another species investigated this way ^51^. Songs typically consist of hierarchically structured, meaningless sound elements and variations in their composition rarely alter the informational content (see Engesser & Townsend, 2019 for review). In contrast, sooty mangabey vocal sequences are built from calls that can be emitted individually or combined into short sequences of two call types, or longer iterative sequences containing the same two call types (e.g., grunt_twitter), which follow hierarchical rules resulting in predictable structures. Such a system is constrained in that there are limitations in the call types that can combine together into an utterance. It nonetheless provides several different predictable structures to which meaning could potentially be bound. Thus, such a system holds the capacity to expand messaging well beyond the number of utterances available if two calls combined in only one way and suggests that large repertoires of distinct sequences are not the only option for generating a wide variety of utterances. Whether these predictable sequences carry different meanings remains to be tested.

**Table 4.** Hierarchical structures in sooty mangabey common vocal sequences G represents a ‘grunt’ call, T a ‘twitter’ call. + refers to the call type in the sequence with the highest occurrence of element repetitions. (GT)n means any number of GT. (TG)n means any number of TG.

| Sequence type | N sequence | N individual | Hierarchical structure | Explanation |
| --- | --- | --- | --- | --- |
| GTG+ | 44 | 16 | Recombination<br>Non-adjacent dependencies | GT is recombined with a final G+<br>First G calls for G+ at the end |
| GT+GTG+ | 15 | 7 | Recombination<br>Non-adjacent dependencies | GT+ and GT are recombined together, and end with a final G+<br>First G calls for G+ at the end |
| GT+(GT)<sup>n</sup>GTG+ | 7 | 4 | Recombination<br>Non-adjacent dependencies<br>Recursion | GT+ and GT are recombined together and with a final G+<br>First G calls for G+ at the end<br>GT is embedded and recursively occurs several times in the sequence. |
| T+GT<br>T+GTG+<br>T+GTG+T | 8<br>9<br>2 | 4<br>6<br>2 | Recombination | T+G recombined with a final T, a TG+ or another GT |
| T+(GT)<sup>n</sup>GTG+<br>T+(GT)<sup>n</sup>GTG+T | 8<br>3 | 6<br>3 | Recombination<br>Recursion | T+G recombined with a TG+ or another GT<br>GT recombined and also embedded and recursively occurs several times in the sequence |

### Sequence flexibility

Sooty mangabeys primarily produced single calls (82.9% of female vocal production and 75.9% of male vocal production) and both sexes can combine most of their calls (nine out of 10 call types, respectively) into sequences. However, some call types—’hoo’, ‘grumble’, and ‘wau’—were almost exclusively produced in sequences. Calls restricted to sequences occur widely across taxa though functional explanations beyond affixation remain unclear, e.g., apes 48, bats 19, birds 20,24,27–29, cetaceans 66–69, frogs 7, mongooses 55, monkeys ^37–39,42,43,45,49,70–74^.

Without iterations, sooty mangabeys produced sequences of two to four distinct call types, consistent with patterns observed across many species e.g., apes ^12–14,16–18,51^, birds ^20,21,23,24,28,29,75–78^, cetaceans ^68,69^, elephants ^8,79,80^, frogs (Bhat et al., 2022; Deng et al., 2024), mongooses ^30,55,83^, monkeys ^32,33,36–40,42–46,49,70–72,84–91^, although most sequences comprised two calls. Longer sequences combining six to eight different call types are less frequently documented but have been reported in Amboli bush frogs (Bhat et al., 2022), geladas ^92^, greater horseshoe bats ^19^, meerkats ^31^, and chimpanzees ^48^. When iterations are included, sooty mangabeys showed sequence lengths comparable to those of chimpanzees (up to 10 call types) and marmosets (up to nine call types) ^48,52^.

The frequencies of individual call types used in sequences, the types of combination pairs and unique sequences, were unevenly distributed (Figure 1, Figure S1, Tables S1, S2, S3 & S4).

All call types were found in sequences produced by at least eight individuals with a minimum of 20 occurrences, except ‘waus’ that were found in four sequences produced by one individual, ‘vibratos’ and ‘whoop-gobbles’ that were only produced singly. Females predominantly used ‘grunts’ and ‘twitters’ (79.5% of recorded calls in sequences), while males primarily used ‘shrills’ and ‘hoos’ (78.2% of recorded calls in sequences) (Figure 1). Five sequences with iterations were predominant, requiring only four call types: grunt_twitter, twitter_grunt and grunt_twitter_grunt in females and shrill_hoo and shrill_hoo_shrill_hoo in males. Notably, more than half of the unique sequences with iterations occurred only once in the dataset (21/41 in females and 14/21 in males) (Tables S2 & S4). To facilitate comparison across species, we focus on two metrics contributing to vocal system flexibility and that are the most frequently reported in the literature: the overall number of call types that are combined and the number of distinct sequences.

Sooty mangabeys—who can combine eight out of their 10 call types—are at the upper end of the range of call type usage in sequences documented in animals, alongside chimpanzees ^15,48^, common marmosets ^52^, black-and-white ruffed lemurs ^84^, Japanese great tits ^28^, cotton- top tamarins ^86^ and orcas ^68^. This is higher than species like Campbell’s monkeys ^72^ and Philippine tarsiers ^90^ which combine about half of their call types, as well as De Brazza’s monkeys ^71^, Sahamalaza sportive lemurs ^89^, and mongoose lemurs ^74^, which combine less than a third of their call types. Surprisingly, closely related red-capped mangabeys combine fewer of their calls (seven out of 12) ^70^, although this study was conducted in captivity where social and environmental factors may influence call and sequence use.

The second metric useful for comparative purposes, the number of possible sequences (excluding iterations), shows that sooty mangabeys produced 28 different sequences. Compared to species with a similar vocal repertoire size, sooty mangabeys exhibit greater sequence diversity than Japanese great tits with 19 reported sequences ^28^, De Brazza’s monkeys—1—^71^, Campbell’s monkeys—3—^72^, red-capped mangabeys—8—^70^ and sahamalaza sportive lemurs—1—^89^, though they fall well below chimpanzees—282—^48^and marmosets—7 different bigrams and 56 different trigrams reported—^52^. While sooty mangabeys can produce a considerable variety of sequences and are flexible in the number of different call types they can combine, the majority of their sequences use few call types overall, focusing mainly on sequences of two call types. This suggests they primarily use a limited set of common sequences, while other sequences occur less frequently but remain noteworthy.

Despite their ability to combine most calls, sooty mangabeys do not do so frequently, which requires explanation. In Shannon’s theory of information, systems with repetitive use of few signals have low information entropy, conveying less information compared to systems with greater signal diversity ^93^. Sooty mangabeys might also use less common, unstructured sequences to convey meanings. Less common sequences often involve the same calls but arranged in different orders, specifically ‘scream’, ‘hoo’, ‘grumble’ and ‘growl’ usually emitted in aggression contexts^57^ (Tables S1, S2, S3 & S4). These sequences could result from call production during rapid changes of events ^94^ (e.g., in aggressive contexts where social exchanges can unfold at speed with different levels of aggressive behaviours or bystander individuals joining the fight), offering a read-out of rapid social shifts to non-visible conspecifics. However, confirming this hypothesis would require further investigation into the contexts of sequence production. Another possibility is under-sampling of these sequences, particularly in contexts that pose challenges for data collection (see limitations).

### Sex differences

Overall, females but never males produced sequences composed of ‘grunts’ and ‘twitters’ (note that adult males do not produce ‘twitters’). This aligns with findings in closely related red-capped mangabeys where females produce more sequence types and at higher rates than males, likely because their social role is to mediate intragroup social relationships ^70^. This contrasts with chimpanzees, which share vocal sequences across sexes ^95^, suggesting usage constraints in *catarrhine* monkeys compared to apes

### Limitations

Despite careful observations, some vocal utterances in some contexts were difficult to capture and hence may be underrepresented in the dataset, like events occurring in tall trees, at night, or during rapidly unfolding aggression contexts where vocalisations from different signallers were often simultaneous. Moreover, call types were classified into broad categories, which may overlook finer acoustic variations within each type. For instance, ‘twitters’ are suggested to have acoustic variations depending on the context of production ^57^. If such variations carry specific meaning, they could impact sequence structures, which we did not consider in our analyses.

## Conclusion

Sooty mangabeys produce rule-based sequences that include hierarchical structures which have been rarely documented in animal communication outside of songs. They also combine some calls without using specific structure, with half of their sequence types recorded being idiosyncratic. These mechanisms are likely promoted by different contexts. An advantage of different rule-based sequences using the same call types, such a twitter-grunt and grunt- twitter, may be to convey different meanings depending on the order of production, or on non-adjacent dependencies across extended sequences. Such sequences would for instance allow unambiguous messaging even in low-visibility dense forest habitats in which sooty mangabeys live. Indeed, ‘grunts’ and ‘twitters’ are produced in a wide range of contexts, from daily activities such as moving and feeding to affiliative behaviours ^57,96^. This contextual variation will allow future research to determine whether emitting these calls either singly and in differently structured sequences conveys distinct meanings to receivers. In contrast, unstructured sequences may be promoted by rapid changes of events, offering for instance a flexibly produced readout of the unfolding of rapid social shifts during aggressive contexts. These findings highlight the importance of a whole-repertoire approach to assess sequence usage and associated rules, as this is key to understanding how vocal systems can potentially expand meanings beyond the number of vocalisations. Critically, sooty mangabeys produce hierarchically structured sequences using only two call types, a system in animal vocal communication that, to our knowledge, has not been previously documented.

## Acknowledgments

We thank the Ministère des Eaux et Forêts and the Office Ivoirien des Parcs et Réserves in Ivory Coast for permitting this study (Research permit: Wittig/006/MESRS/DRI). Special thanks to Christophe Boesch for establishing and nurturing the Taï Chimpanzee Project, Anderson Bitty, the Centre Suisse de Recherches Scientifiques, field assistants, and all members of the Taï Chimpanzee and Taï Monkey Projects for their invaluable support. We are grateful to Derry Taylor for insightful discussions and to Tatiana Bortolato, Ryan Sigmundson, and Mathilde Grampp for their assistance with methodological design.

