## Supplementary material for "Rule-based sequences in sooty mangabey vocal communication"

### Electronic supplementary material

**Document S1. Call types**

Range and Fischer (2004) have thoroughly described the complete vocal repertoire of sooty mangabeys, and readers seeking more detail should refer to their paper. For convenience, we provide a condensed version here including all the call types examined in our study.

*Growl*

Growls are characterised by a low fundamental frequency band with many visible overtones, and they vary in both frequency and duration, often comprising a combination of shorter and prolonged elements.

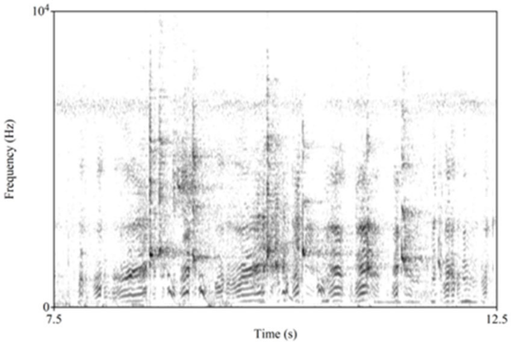


Spectrographic examples of growls. y axis represents the frequency in Hz and x axis the time in s.

*Grumble*

Grumbles are longer than growls, characterised by a low fundamental frequency and a rich harmonic structure with up to 12 overtones.

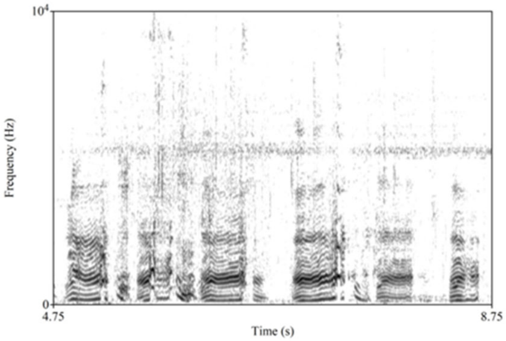


Spectrographic examples of grumbles interspersed with growls (shorter vocalisations). y axis represents the frequency in Hz and x axis the time in s.

*Grunt*

Grunts are low-frequency vocalisations, consistently short in duration (less than 200 milliseconds), and produced across a wide range of contexts. Adult females and males diverge in grunt production (see Range and Fischer).

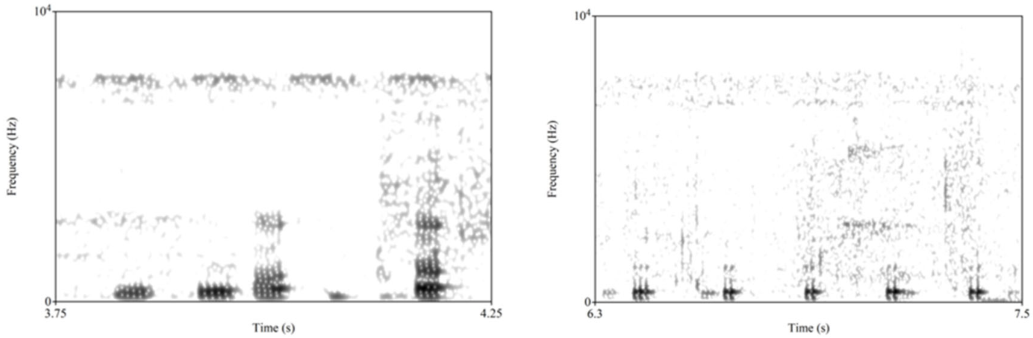


Spectrograms of grunt calls, from an adult female (left) and an adult male (right). y axis represents the frequency in Hz and x axis the time in s.

*Scream*

Screams are harsh calls characterised by wide-band noise extending up to 15 kHz, often lasting for several seconds.

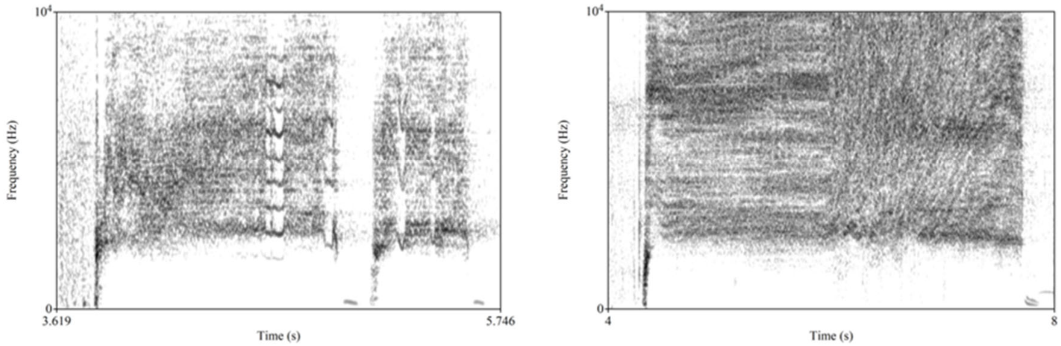


Spectrographic examples of screams. y axis represents the frequency in Hz and x axis the time in s.

*Shrill and hoo* (referred to as alarm call in Range and Fischer, 2004)

Shrill calls consist of several high-frequency elements, sometimes displaying a harmonic structure, though this is less common, and are frequently combined with a hoo call, which features a low fundamental frequency and a rich, readily visible harmonic structure.

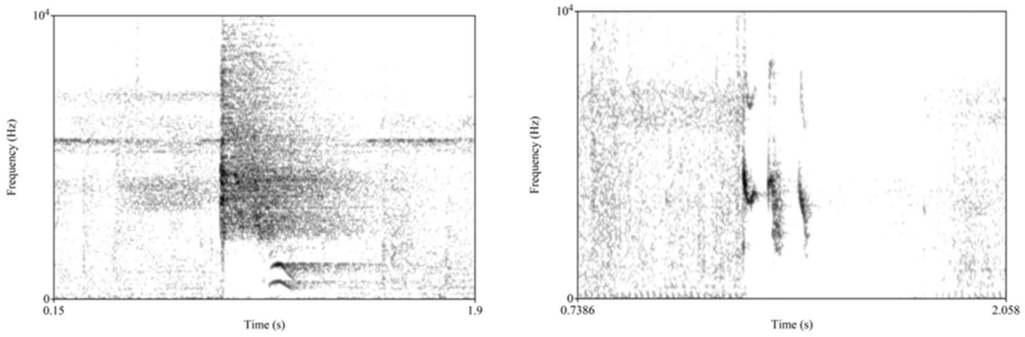


Spectrograms of shrill calls combined with a hoo call (left) and shrill call produced singly (right). y axis represents the frequency in Hz and x axis the time in s.

*Twitter*

Twitters are short, high-frequency calls with a fundamental frequency around 1 kHz and many visible overtones extending up to 20 kHz. Adult males do not produce them.

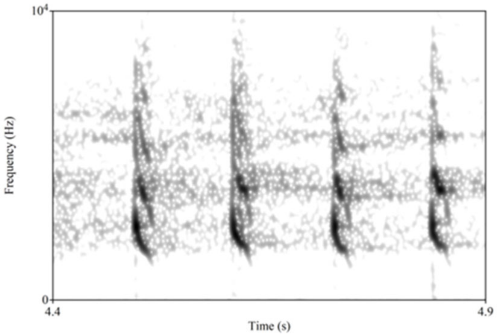


Spectrographic example of twitter. y axis represents the frequency in Hz and x axis the time in s.

*Vibrato* (referred to as copulation call in Range and Fischer, 2004)

Complex call used exclusively by sexually mature females. It consists of many variable elements with a low fundamental frequency and a rich harmonic structure, with individual elements alternating between pitch shifts upwards and downwards, creating the characteristic "vibrato."

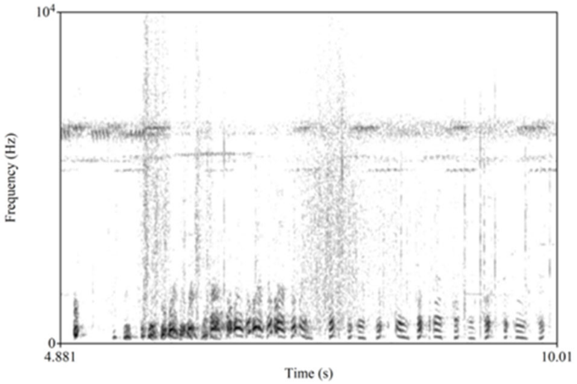


Spectrographic example of vibrato. y axis represents the frequency in Hz and x axis the time in s.

*Wau*

Short, low-frequency call with an ascending-descending pitch pattern and several visible harmonics.

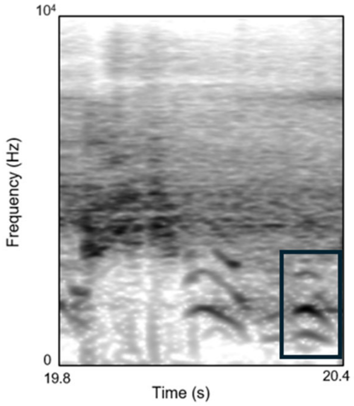


Spectrographic example of wau (in the black box) preceded by shrill and hoo. y axis represents the frequency in Hz and x axis the time in s.

*Whoop-gobble*

Loud, low-frequency complex call that can be heard over long distances and produced by adult males. It begins with a single low-frequency element, followed by a pause that may last several seconds, before concluding with a series of repetitive, low-frequency elements. Although the pause between the first and final elements typically violates our usual rule for defining a call type, we treated it as a special case due to its consistent production in this form.

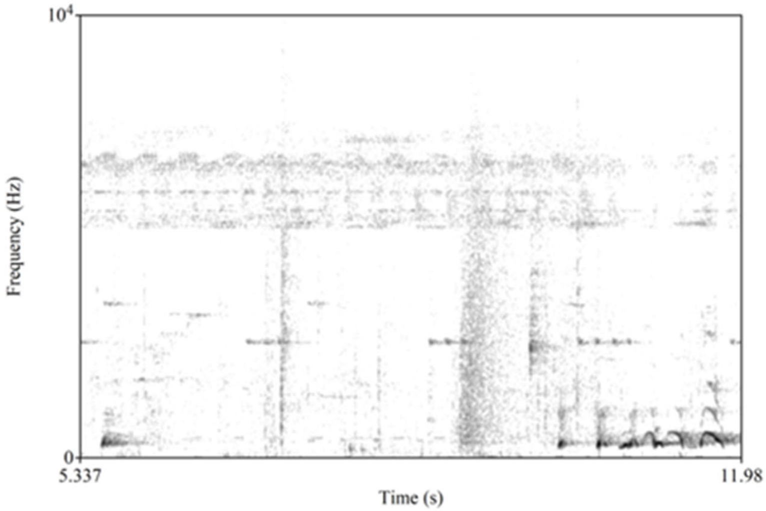


Spectrographic example of whoop-gobble. y axis represents the frequency in Hz and x axis the time in s.

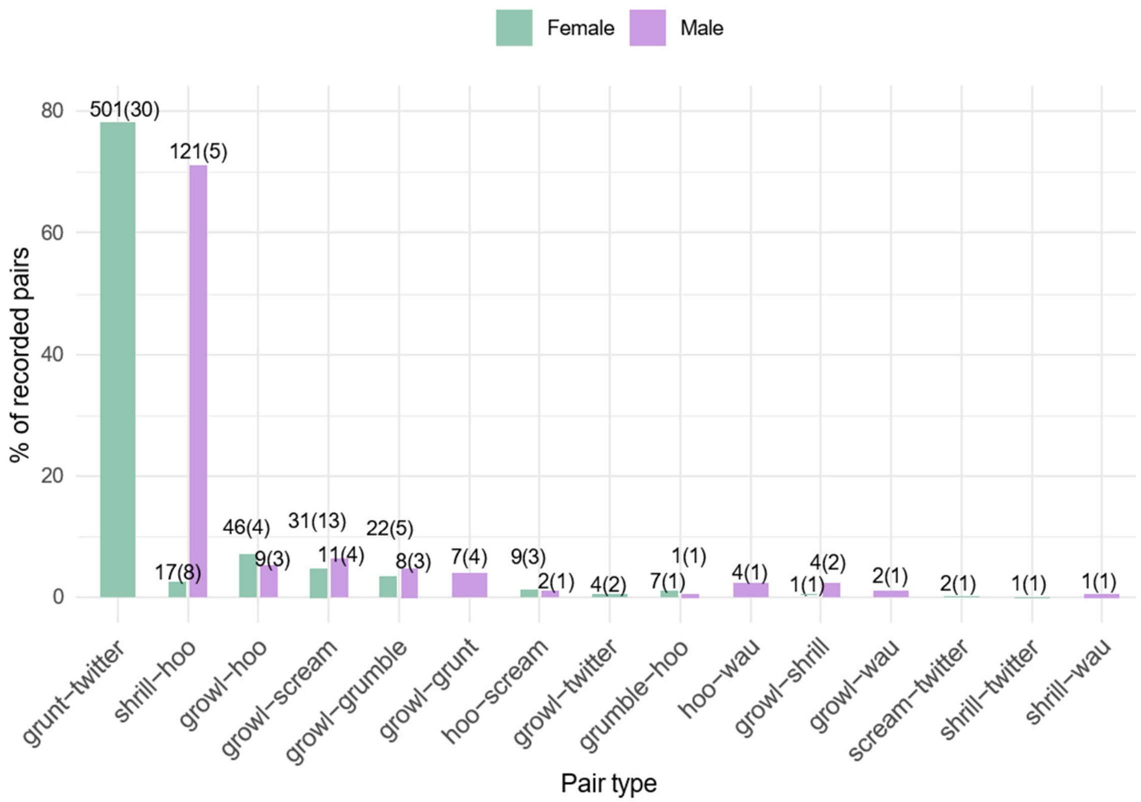


**Figure S1. Percentage of combination pairs recorded per sex**

Above each bar is indicated the sample size with the number of pair recorded next to the number of individuals producing them in brackets.

**Table S1. Female sequence types without call iterations**

| Sequence type | N sequence | N individual | % |
|---|---|---|---|
| grunt_twitter | 153 | 28 | 54.4 |
| twitter_grunt | 74 | 20 | 26.3 |
| shrill_hoo | 16 | 8 | 5.7 |
| growl_scream | 11 | 8 | 3.9 |
| scream_growl | 7 | 7 | 2.5 |
| growl_grumble | 3 | 3 | 1.1 |

| | | | |
|---|---|---|---|
| growl_hoo | 3 | 3 | 1.1 |
| growl_hoo_grumble | 3 | 1 | 1.1 |
| scream_hoo | 3 | 2 | 1.1 |
| growl_scream_grumble | 1 | 1 | 0.4 |
| growl_scream_hoo | 1 | 1 | 0.4 |
| grumble_growl | 1 | 1 | 0.4 |
| grunt_twitter_growl | 1 | 1 | 0.4 |
| scream_growl_grumble | 1 | 1 | 0.4 |
| scream_twitter_growl | 1 | 1 | 0.4 |
| twitter_growl | 1 | 1 | 0.4 |
| twitter_growl_shrill | 1 | 1 | 0.4 |

**Table S2. Female sequence types with call iterations**

| **Sequence type** | **N sequence** | **N individual** | **%** |
|---|---|---|---|
| grunt_twitter | 82 | 26 | 29.2 |
| grunt_twitter_grunt | 44 | 16 | 15.7 |
| twitter_grunt | 44 | 17 | 15.7 |
| grunt_twitter_grunt_twitter_grunt | 15 | 7 | 5.3 |
| shrill_hoo | 15 | 7 | 5.3 |
| growl_scream | 9 | 6 | 3.2 |

| | | | |
|---|---|---|---|
| twitter_grunt_twitter_grunt | 9 | 6 | 3.2 |
| twitter_grunt_twitter | 8 | 4 | 2.8 |
| grunt_twitter_grunt_twitter_grunt_twitter_grunt | 5 | 3 | 1.8 |
| scream_growl | 4 | 4 | 1.4 |
| twitter_grunt_twitter_grunt_twitter_grunt_twitter_grunt | 4 | 4 | 1.4 |
| growl_grumble_growl | 3 | 3 | 1.1 |
| grunt_twitter_grunt_twitter | 3 | 2 | 1.1 |
| scream_growl_scream | 3 | 3 | 1.1 |
| growl_hoo_growl_hoo_growl | 2 | 2 | 0.7 |
| growl_scream_growl | 2 | 2 | 0.7 |
| grunt_twitter_grunt_twitter_grunt_twitter | 2 | 2 | 0.7 |
| grunt_twitter_grunt_twitter_grunt_twitter_grunt_twitter_grunt | 2 | 2 | 0.7 |
| twitter_grunt_twitter_grunt_twitter | 2 | 2 | 0.7 |
| twitter_grunt_twitter_grunt_twitter_grunt | 2 | 2 | 0.7 |
| growl_hoo_growl_grumble_growl | 1 | 1 | 0.4 |
| growl_hoo_growl_hoo_growl_hoo_growl | 1 | 1 | 0.4 |
| growl_hoo_growl_hoo_growl_hoo_growl_grumble | 1 | 1 | 0.4 |
| growl_hoo_growl_hoo_growl_hoo_growl_hoo_growl_hoo_growl_hoo_grumble_hoo_grumble_growl_grumble_hoo_growl_hoo_grumble_growl_hoo_growl_hoo_grumble_hoo_growl_grumble_growl_hoo_growl_grumble_growl_hoo_growl_hoo_growl | 1 | 1 | 0.4 |

| | | | |
|---|---|---|---|
| growl_scream_growl_grumble | 1 | 1 | 0.4 |
| growl_scream_growl_scream_hoo_growl_scream_hoo | 1 | 1 | 0.4 |
| grumble_growl | 1 | 1 | 0.4 |
| grunt_twitter_grunt_twitter_grunt_twitter_growl | 1 | 1 | 0.4 |
| scream_growl_grumble_growl_grumble_growl | 1 | 1 | 0.4 |
| scream_hoo | 1 | 1 | 0.4 |
| scream_hoo_scream | 1 | 1 | 0.4 |
| scream_hoo_scream_hoo_scream | 1 | 1 | 0.4 |
| scream_twitter_scream_growl | 1 | 1 | 0.4 |
| shrill_hoo_shrill | 1 | 1 | 0.4 |
| twitter_growl | 1 | 1 | 0.4 |
| twitter_growl_shrill_twitter_growl | 1 | 1 | 0.4 |
| twitter_grunt_twitter_grunt_twitter_grunt_twitter | 1 | 1 | 0.4 |
| twitter_grunt_twitter_grunt_twitter_grunt_twitter_grunt_twitter | 1 | 1 | 0.4 |
| twitter_grunt_twitter_grunt_twitter_grunt_twitter_grunt_twitter_grunt | 1 | 1 | 0.4 |
| twitter_grunt_twitter_grunt_twitter_grunt_twitter_grunt_twitter_grunt_twitter_grunt_twitter_grunt_twitter_grunt | 1 | 1 | 0.4 |
| twitter_grunt_twitter_grunt_twitter_grunt_twitter_grunt_twitter_grunt_twitter_grunt_twitter_grunt_twitter_grunt_twitter_grunt_twitter_grunt_twitter_grunt_twitter_grunt_twitter_grunt_twitter_grunt_twitter | 1 | 1 | 0.4 |

**Table S3. Male sequence types without call iterations**

| Sequence type | N sequence | N individual | % |
|---|---|---|---|
| shrill_hoo | 66 | 5 | 72.5 |
| shrill_hoo_growl | 6 | 1 | 6.6 |
| growl_scream | 5 | 4 | 5.5 |
| growl_shrill_hoo | 3 | 2 | 3.3 |
| grunt_growl | 3 | 2 | 3.3 |
| shrill_hoo_wau | 2 | 1 | 2.2 |
| shrill_hoo_wau_growl | 2 | 1 | 2.2 |
| growl_grumble_grunt | 1 | 1 | 1.1 |
| growl_scream_hoo | 1 | 1 | 1.1 |
| grunt_growl_grumble | 1 | 1 | 1.1 |
| scream_hoo_grumble_growl | 1 | 1 | 1.1 |

**Table S4. Male sequence types with call iterations**

| Sequence type | N sequence | N individual | % |
|---|---|---|---|
| shrill_hoo | 49 | 4 | 53.8 |
| shrill_hoo_shrill_hoo | 16 | 4 | 17.6 |
| shrill_hoo_growl | 4 | 1 | 4.4 |
| growl_scream | 2 | 2 | 2.2 |

| | | | |
|---|---|---|---|
| growl_scream_growl | 2 | 2 | 2.2 |
| growl_shrill_hoo_growl | 2 | 2 | 2.2 |
| grunt_growl | 2 | 1 | 2.2 |
| growl_grumble_growl_grunt | 1 | 1 | 1.1 |
| growl_scream_growl_scream_growl | 1 | 1 | 1.1 |
| growl_scream_hoo_growl | 1 | 1 | 1.1 |
| growl_shrill_hoo | 1 | 1 | 1.1 |
| grunt_growl_grumble_growl_grumble | 1 | 1 | 1.1 |
| grunt_growl_grunt_growl | 1 | 1 | 1.1 |
| scream_hoo_grumble_growl_grumble_growl | 1 | 1 | 1.1 |
| shrill_hoo_growl_shrill_hoo | 1 | 1 | 1.1 |
| shrill_hoo_shrill_hoo_growl | 1 | 1 | 1.1 |
| shrill_hoo_shrill_hoo_shrill_hoo | 1 | 1 | 1.1 |
| shrill_hoo_shrill_hoo_wau_growl | 1 | 1 | 1.1 |
| shrill_hoo_wau | 1 | 1 | 1.1 |
| shrill_hoo_wau_growl | 1 | 1 | 1.1 |
| shrill_hoo_wau_shrill_hoo | 1 | 1 | 1.1 |

**Table S5. Result models of positional bias analysis**

| **Parameters** | **Estimate** | **Error** | **Lower 95%CI** | **Upper 95%CI** | **Bulk ESS** | **Tail ESS** |
|---|---|---|---|---|---|---|

| | | | | | | |
|---|---|---|---|---|---|---|
| Model - Growl | | | | | | |
| Intercept | -0.38 | 0.44 | -1.27 | 0.45 | 6426 | 4171 |
| group | -0.8 | 0.83 | -2.53 | 0.78 | 4711 | 4004 |
| Model - Grunt | | | | | | |
| sd(Intercept) - identity | 0.98 | 0.34 | 0.44 | 1.73 | 3703 | 4269 |
| Intercept | -0.66 | 0.31 | -1.34 | -0.07 | 3703 | 4269 |
| group | -1.25 | 0.65 | -2.55 | -0.01 | 5039 | 5108 |
| Model - Scream | | | | | | |
| Intercept | 0.25 | 0.52 | -0.77 | 1.28 | 5973 | 5128 |
| group | 0.68 | 0.9 | -1.01 | 2.56 | 5750 | 4681 |
| Model - Twitter | | | | | | |
| sd(Intercept) - identity | 0.95 | 0.33 | 0.42 | 1.72 | 2472 | 3140 |
| Intercept | 0.56 | 0.31 | -0.03 | 1.2 | 3753 | 5105 |
| group | 1.33 | 0.66 | 0.11 | 2.7 | 5554 | 5341 |

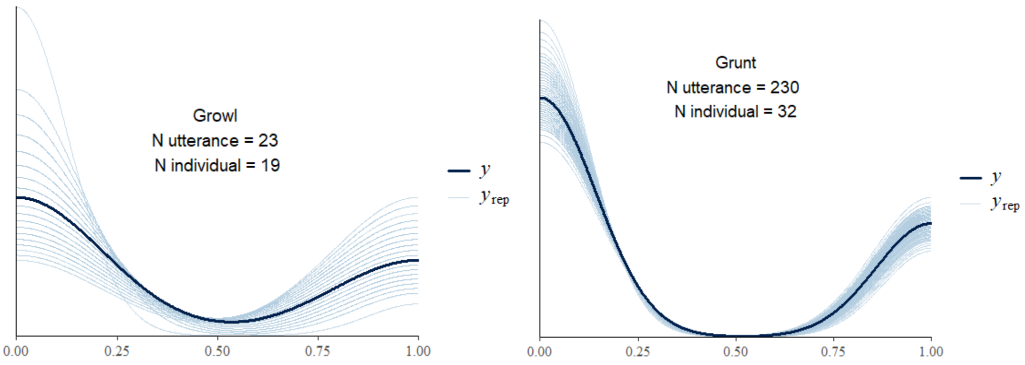

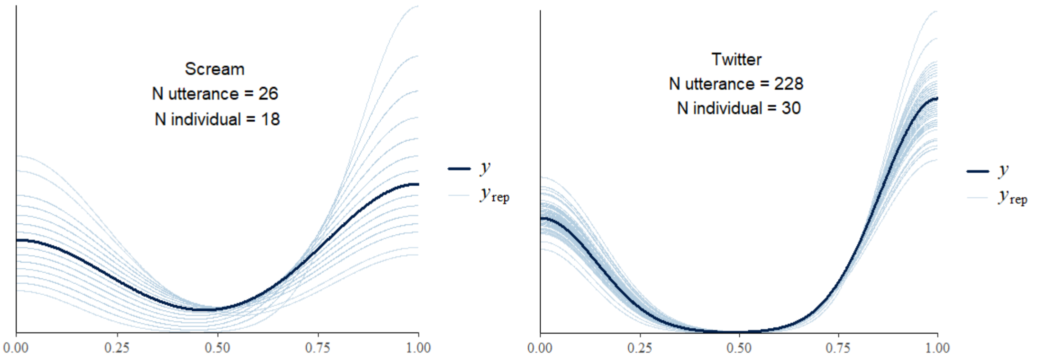


**Figure S2. Posterior predictive of models for positional bias analysis**

**Table S6. Result models of call iteration analysis (grunt and twitter sequences)**

| **Parameters** | **Estimate** | **Error** | **Lower 95%CI** | **Upper 95%CI** | **Bulk ESS** | **Tail ESS** |
|---|---|---|---|---|---|---|
| Model - Grunt_Twitter+ | | | | | | |
| sd(Intercept) - identity | 2.64 | 1.67 | 0.32 | 6.99 | 1833 | 1876 |
| Intercept | -4.53 | 1.92 | -9.51 | -2.02 | 2480 | 1334 |
| group | 1 | 2.45 | -3.77 | 6.24 | 2809 | 2605 |
| Model - Twitter_Grunt+ | | | | | | |
| sd(Intercept) - identity | 2.16 | 1.43 | 0.22 | 5.8 | 2222 | 2168 |
| Intercept | -0.72 | 0.92 | -2.84 | 0.93 | 2863 | 2489 |

*Posterior predictive*

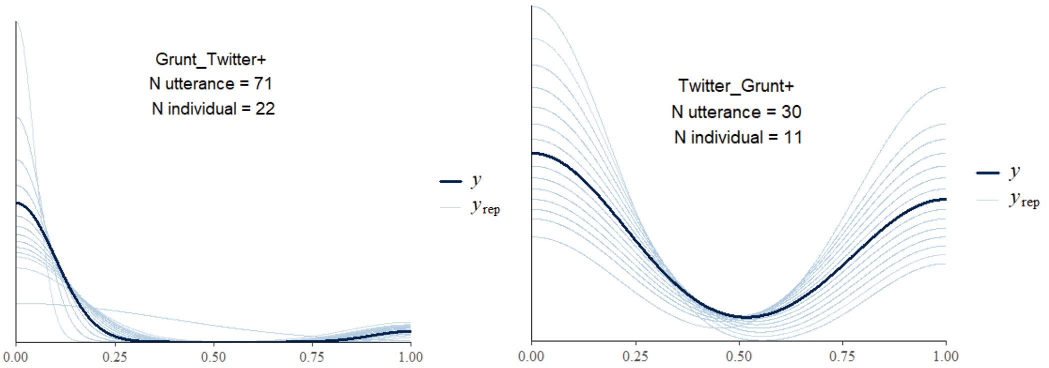


**Figure S3. Posterior predictive of models for call iteration analysis**

**Table S7. Result models of element repetition analysis (grunt and twitter sequences)**

| **Parameters** | **Estimate** | **Est.Error** | **l-95% CI** | **u-95% CI** | **Bulk_ESS** | **Tail_ESS** |
|---|---|---|---|---|---|---|
| Model - Grunt_Twitter - Grunt elements | | | | | | |
| Intercept | 0.87 | 0.14 | 0.59 | 1.16 | 2659.54 | 3904.81 |
| Last position | 1.03 | 0.09 | 0.86 | 1.20 | 5035.49 | 5684.21 |
| Middle position | 0.21 | 0.14 | -0.07 | 0.49 | 5780.39 | 6260.90 |
| Length 3 | 0.07 | 0.12 | -0.17 | 0.30 | 3978.21 | 5214.52 |
| Length 5 | 0.01 | 0.14 | -0.27 | 0.28 | 3989.47 | 5229.73 |
| Length 7 | 0.34 | 0.17 | 0.01 | 0.66 | 4418.67 | 6032.49 |
| Group | -0.09 | 0.23 | -0.55 | 0.37 | 2402.11 | 3379.56 |
| sd(Intercept) | 0.49 | 0.09 | 0.34 | 0.71 | 2365.29 | 3784.06 |
| Model - Grunt_Twitter - Twitter elements | | | | | | |
| Intercept | 1.66 | 0.09 | 1.47 | 1.83 | 5195.15 | 5034.86 |
| Last position | -0.72 | 0.16 | -1.05 | -0.40 | 12992.66 | 6581.66 |

| | | | | | | |
|---|---|---|---|---|---|---|
| Middle position | -0.89 | 0.41 | -1.74 | -0.11 | 11916.51 | 6302.61 |
| Length 3 | -0.03 | 0.09 | -0.21 | 0.15 | 11869.33 | 6100.93 |
| Length 5 | 0.17 | 0.12 | -0.07 | 0.41 | 10335.92 | 6562.52 |
| Length 7 | -0.25 | 0.22 | -0.68 | 0.16 | 10425.66 | 6343.10 |
| Group | -0.02 | 0.14 | -0.30 | 0.27 | 6093.67 | 5872.45 |
| sd(Intercept) | 0.26 | 0.07 | 0.15 | 0.42 | 2852.71 | 4629.38 |
| Model - Twitter_Grunt - Grunt elements | | | | | | |
| Intercept | 1.26 | 0.22 | 0.81 | 1.68 | 2377.77 | 3699.90 |
| Last position | 1.20 | 0.26 | 0.72 | 1.72 | 7323.14 | 5471.76 |
| Length 3 | -0.62 | 0.31 | -1.26 | -0.04 | 7715.19 | 5720.03 |
| Length 4 | -0.68 | 0.26 | -1.21 | -0.17 | 6760.46 | 5665.66 |
| Group | 1.06 | 0.43 | 0.23 | 1.91 | 2911.14 | 3956.91 |
| sd(Intercept) | 0.74 | 0.16 | 0.49 | 1.12 | 2680.45 | 4101.65 |
| Model - Twitter_Grunt - Twitter elements | | | | | | |
| Intercept | 2.00 | 0.13 | 1.74 | 2.24 | 2997.06 | 4325.24 |
| Last position | -1.06 | 0.16 | -1.39 | -0.74 | 10989.22 | 5845.54 |
| Length 3 | 0.29 | 0.16 | -0.03 | 0.61 | 6764.18 | 5575.56 |
| Length 4 | 0.09 | 0.15 | -0.20 | 0.38 | 8353.47 | 5932.06 |
| Group | -0.30 | 0.28 | -0.85 | 0.26 | 4557.00 | 5043.04 |
| sd(Intercept) | 0.39 | 0.10 | 0.23 | 0.64 | 3031.85 | 4861.99 |

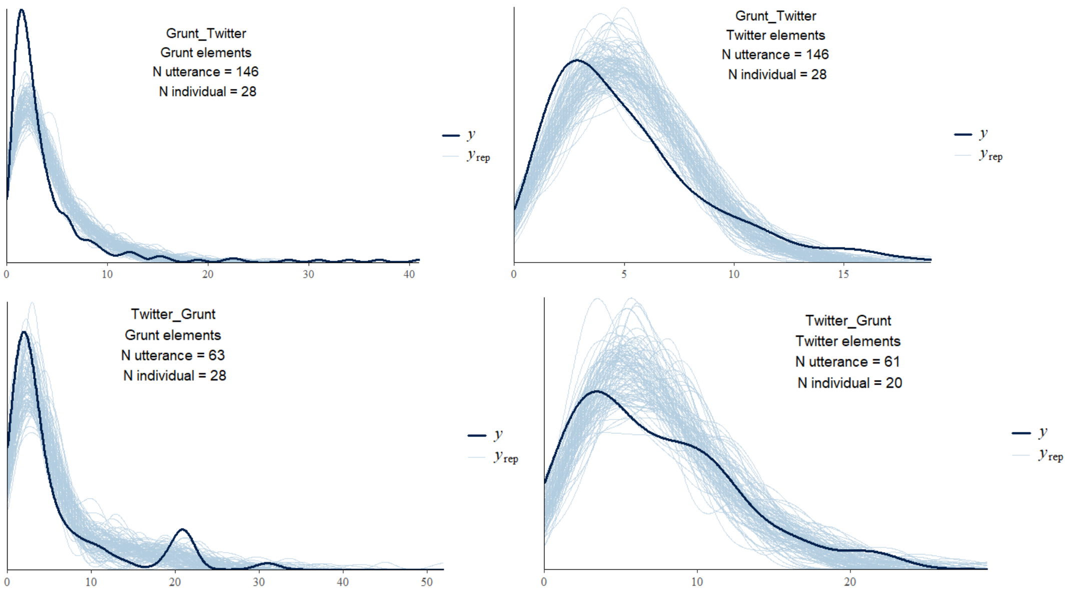


**Figure S4. Posterior predictive of models for element repetition analysis**

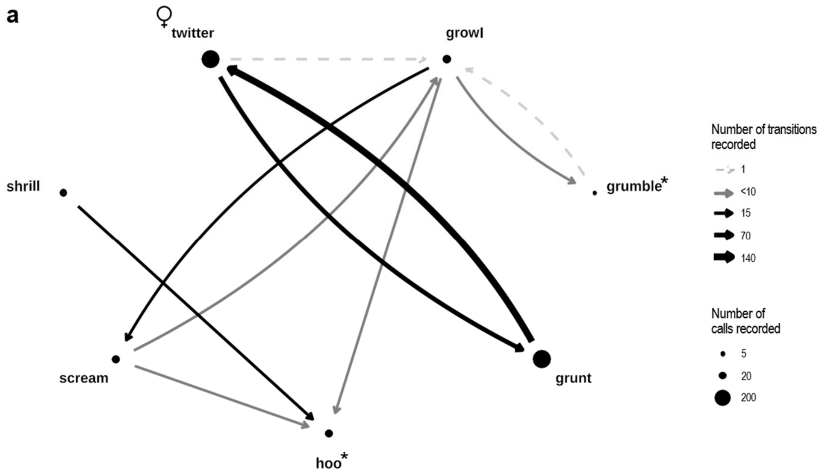

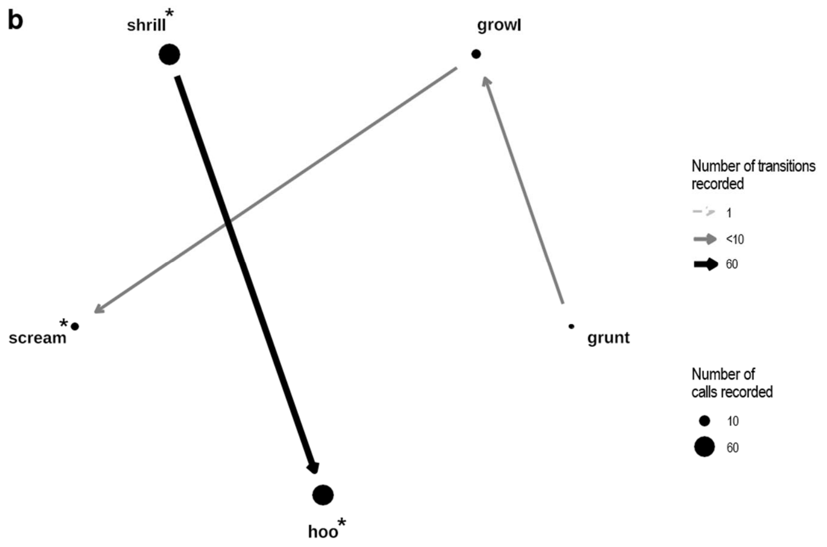


**Figure S5. 2-call type sequences without iterations of adult sooty mangabey a females and b males**

These networks represent all recorded call types and transitions between the different call types in sequences without iterations. * = call type produced exclusively in combination with another call type (i.e., never or idiosyncratic -maximum one occurrence- singly). ♀ = call type produced only by females. Node (circle) size is proportional to the occurrence of call for each call type and edge: (line) thickness and color are proportional to the number of combinations.
